# Computational design of improved standardized chemotherapy protocols for grade II oligodendrogliomas

**DOI:** 10.1101/521559

**Authors:** Víctor M. Pérez-García, Luis E. Ayala-Hernández, Juan Belmonte-Beitia, Philippe Schucht, Michael Murek, Andreas Raabe, Juan Sepúlveda

## Abstract

The use of mathematical models for personalization of cancer therapies and raising hypothesis of potential clinical impact is an emerging topic in the interface between mathematics and oncology. Here we put forward a mathematical model describing the response of low-grade (WHO grade II) oligodendrogliomas (LGO) to temozolomide (TMZ). The model described the longitudinal volumetric dynamics of tumor response to TMZ of a cohort of 11 LGO patients treated with TMZ. After finding patient-specific parameters, different therapeutical strategies were tried computationally on the ‘in-silico twins’ of those patients. Chemotherapy schedules with larger-than-standard rest periods between consecutive cycles had either the same or better long-term efficacy than the standard 28-day cycles. The results were confirmed in a large virtual clinical trial including 2000 patients. These long-cycle schemes would also have reduced toxicity and defer the appearance of resistances.

On the basis of those results, a combination scheme consisting of five induction TMZ cycles given monthly plus 12 maintenance cycles given every three months was found to provide substantial survival benefits for the in-silico twins of the 11 LGO patients (median 5.69 years, range: 0.67 to 68.45 years) and in a large virtual trial including 2000 patients. This scheme could be useful for defining a standardized TMZ treatment for LGO patients with survival benefits.

**Author summary:** A mathematical model described the longitudinal volumetric growth data of grade II oligodendrogliomas patients and their response to temozolomide. The model was used to explore alternative therapeutical protocols for the in-silico twins of the patients and in virtual clinical trials. The simulations show that enlarging the time interval between chemotherapy cycles would maintain the therapeutical efficacy, while limiting toxicity and defering the development of resistances. This may allow for improved drug-exposure by administering a larger number of cycles for longer treatment periods. A scheme based on this idea consisting of an induction phase (5 consecutive cycles, 1 per month) and a maintenance phase (12 cycles given in three-months intervals) led to substantial survival benefits in-silico.

## Introduction

Oligodendrogliomas (ODGs) are low-incidence glial tumors, affecting mostly young adults. They are slowly growing, infiltrative tumors with isocitrate dehydrogenase 1 or 2 mutations and codeletion of chromosomal arms 1p and 19q. Grade II ODGs (LGO) are well differentiated tumors with a low mitotic index [1]. In spite of the long median patient survival, they are incurable currently [2].

Many ODG patients present few neurological symptoms for extended periods of time. The decision on the specific combination of therapies to be used on each patient is based on the qualitative consideration of different variables including age, tumor grade, performance status and tumor location [3]. Radiation therapy (RT) is beneficial for patients in terms of survival, but its timing has been the subject of debate [4]. Regarding chemotherapy (CT), temozolomide (TMZ), an oral alkylating agent, has a favourable toxicity profile [5] and can contribute to reduction in seizure frequency in low-grade glioma (LGG) patients [6]. Phase II trials have demonstrated its effectivity against LGGs [7–9]. Also, neoadjuvant CT given to surgically unresectable tumors has allowed subsequent gross total resection in some cases [10], which is of relevance when the tumour is highly infiltrative or located in eloquent areas. Thus, prolonged TMZ treatment is a relevant option either as up-front or as adjuvant treatment.

Clinical trials have shown a similar efficacy of TMZ vs RT for 1p/19q-codeleted tumors [11, 12]. Also, RT is associated with late neurocognitive toxicity. Thus CT is frequently used as first-line treatment for ODG patients. In that context, relevant questions arise such choice of the chemotherapy regimen and the optimal number of cycles to be prescribed.

Mathematical models have potential to help in finding optimized treatment schedules/combinations improving survival and/or reducing toxicity [13, 14]. Once the base mathematical model is set, patient-specific parameters can be obtained from data. That provides an ‘in-silico twin’ [15] allowing computational studies that could be beneficial for real patients.

## Materials and methods

### Patients

82 patients diagnosed of LGG (biopsy/surgery confirmed astrocytoma, oligoastrocytoma or oligodendroglioma according to the WHO 2007 classification) and followed at the Bern University Hospital between 1990 and 2013 were initially included in the study. The study was approved by Kantonale Ethikkommission Bern (Bern, Switzerland), with approval number: 07.09.72.

Of that patient population, we selected 1p/19q-codeleted tumors (thus LGOs according to the 2016 WHO classification) treated with at least three cycles of TMZ, having no previous RT treatment and no other treatment given in the period of study. Only 16 patients satisfied these criteria. Of them 3 (19%) did not respond to TMZ and 2 (12%) responded initially but progressed during treatment to anaplastic forms. Thus 11 (69%) oligodendroglioma patients responded to the therapy, did not display any signs of malignant transformation and were used for this study.

### Image acquisition and analysis

Radiological glioma growth was quantified by manual measurements of tumour diameters on successive MRIs (T2/FLAIR sequences). Since some of the older patients were available only as jpeg images we computed the volume using the ellipsoidal approximation. The three largest tumour diameters (*D*_1_, *D*_2_, *D*_3_) along the axial, coronal and sagittal planes were measured and tumour volumes estimated using the equation *V* = (*D*_1_·*D*_2_·*D*_3_)/2, following the standard practice [16]. To estimate the error of the methodology we took a different set of glioma patients from another study [17] and compared their volumes computed accurately using a semi-automatic segmentation approach with those computed using the ellipsoidal approximation. Mean differences were 18%, that was the reference level used for the error in the volume computations.

### Mathematical model

In this paper we considered LGOs in a simplified way as composed of two tumor cell compartment. The first one was the tumor cell population *P*(*t*), assumed to grow logistically. The second one was lethally damaged tumor cells because of the action of the therapy *D*(*t*). Temozolomide effect on proliferative cells is a complex one, leading to death through different ‘programmes’ [18–20]. We put together the different processes into two groups, each described by a term in our equations. The first one was early death accounting for necrosis, autophagy and drug-induced apoptosis with rate *α*_1_. The second one was delayed death through mitotic catastrophe with rate *α*_2_. In radiotherapy, the second process is the leading one [21], but not in cytotoxic chemotherapy treatments [19, 20]. The drug concentration in tissue was described by the function *C*(*t*) having a characteristic cleanup time 1/λ.

Figure 1 shows a schematic description of the model. The equations were:

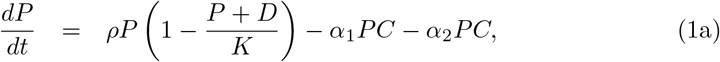

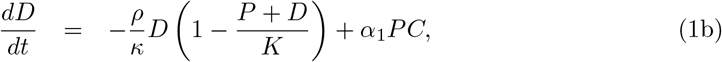

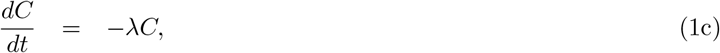

**Fig 1.**
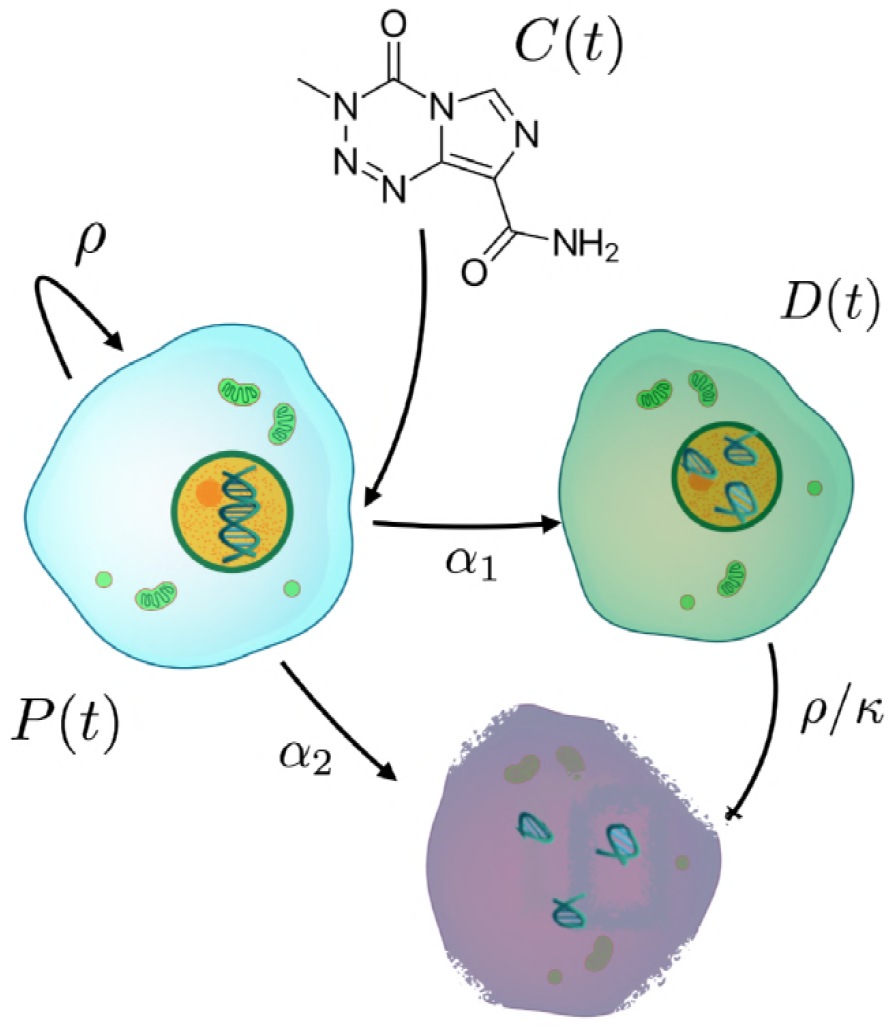
Schematic description of the model defined by Eqs. (1). Tumor cell population *P*(*t*) grows at a rate *ρ* and saturates at a maximum size *K*. These cells are killed by the drug *C*(*t*) (and removed) through direct mechanisms *α*_2_. Another fraction *α*_1_ moves into a different compartment of lethally damaged cells *D*(*t*). These cells die at a rate *ρ/κ* because of mitotic catastrophe.

Chemotherapy was described by a sequence of doses *d*_1_, *d*_2_,…, *d_N_* given at times *t*_1_ < *t*_2_ <…< *t_N_*. The initial time corresponding to the first volumetric observation was denoted as *t*_0_. Initial conditions for Eqs. (1) were taken to be *P*_0_ = *P*(*t*_0_),*D*(*t*_0_) = *C*(*t*_0_) = 0. Drug administration was described as impulses for the times *t_j_* so that 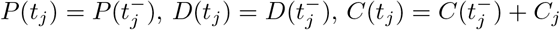 where 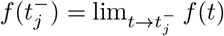 and *C_j_* is the fraction of the dose *d_j_* reaching brain tumor tissue.

### Parameter estimation

We chose the parameter *κ*, corresponding to the averaged number of cell divisions before death by mitotic catastrophe to be equal to 1. The carrying capacity parameter *K* is the one with a less defined value but could be expected to be in a range between 300 and 550 cm^3^. The later number is in line with the maximal volumes observed in LGG patients [22]. However, many patients die when the tumor volume is smaller [15].

The most typical chemotherapy schedule consists of cycles of 28 days with five TMZ oral doses on days 1 to 5 and then a rest period of 23 days. Typical dose per day is *d_j_* = *d* = 150 mg per m^2^ of patient body surface. To calculate the rate of drug decay λ we followed the same methodology as in Ref. [34], using values of TMZ half-life clearance time *t*_1/2_ for doses of 150 mg/m^2^. From the definition of *t*_1/2_ and since Eq. (1c) has exponentially decaying solutions 1/2 = exp (*λt*_1/2_). To account also for the drug loss during transport to the brain we computed the value *C_j_* = *C*_0_ of the dose getting to the tumour as *C_j_* = *β* · *d* · *b*, where *β* is the fraction of TMZ getting to 1 ml of brain interstitial fluid (from a unit dose) and *b* is the patient’s body surface. Then *C_0_* can be interpreted as an effective dose per fraction. The parameter *β* can be calculated using the value of maximal TMZ concentration *C*_max_ for a dose of 150 mg/m^2^ taken from the literature [23]. Since time to reach peak drug concentration in brain is smaller than two hours and thus negligible in comparison with the other time scales in the model, we chose to set the initial drug concentration *C*_0_ to the value *C*_max_ = 0.6 *μ*g/ml as in Ref. [34].

The parameters *α*_1_, *α*_2_ and *ρ* are expected to depend strongly on the tumor growth rate and sensitity to the therapy and will be considered to be adjustable parameters. These parameters, together with the initial population value *P*(0) were fit for each patient longitudinal volumetric data using the library `fmincon` in the scientific software package Matlab (R2017b, The MathWorks, Inc., Natick, MA, USA). Table 1 summarizes the main characteristics and parameter values found for patients included in the study. Since Eqs. (1) are a system of nonlinear ordinary differential equations, it is not possible to find their solutions in closed, explicit, form. Numerical simulations of Eqs. (1) were performed using the Matlab library `ode45`. To test for potential overfitting in the parameter values found, the results for several patients were compared with the results of a brute force approach analyzing a broad parameter range. Both approaches were found to be in excellent agreement. Results for the parameters are listed in Table 1.

**Table 1.** Parameter values best describing the longitudinal volumetric data for the patients included in the study. Values for the other parameters were fixed to *K* = 523.6 cm^3^, λ = 8.3184 day^−1^.

| ID | # CT cycles | $P(0)$ | $\rho$<br>(day<sup>-1</sup>) | $\alpha_1$<br>ml/(μg day) | $\alpha_2$<br>ml/(μg day) |
| --- | --- | --- | --- | --- | --- |
| 6 | 11 | 46.0 | $1.01 \times 10^{-3}$ | 0.32 | 0.1 |
| 10 | 15 | 45.4 | $7.06 \times 10^{-4}$ | 0.27 | 0.21 |
| 25 | 4 | 31.7 | $1.84 \times 10^{-3}$ | 0.76 | 0.75 |
| 57 | 9 | 46.8 | $8.74 \times 10^{-4}$ | 0.23 | 0.21 |
| 105 | 20 | 145.0 | $2.05 \times 10^{-3}$ | 0.1 | 0.14 |
| 108 | 5 | 13.9 | $1.73 \times 10^{-3}$ | 0.92 | 0.18 |
| 151 | 11 | 34.6 | $7.41 \times 10^{-4}$ | 0.52 | 0.05 |
| 159 | 11 | 64.7 | $5.33 \times 10^{-4}$ | 0.57 | 0.6 |
| 170 | 17 | 6.4 | $2.28 \times 10^{-3}$ | 0.01 | 0.28 |
| 203 | 9 | 29.8 | $3.31 \times 10^{-4}$ | 1.99 | 0.1 |
| 213 | 18 | 23.9 | $6.08 \times 10^{-4}$ | 0.3 | 0.3 |

### Virtual clinical trials

To study the effect of the different treatment schedules on patient survival we designed virtual trials. A number of patients was generated by a random choice of the parameters. Uniform distributions were taken for the parameters in the most representative region of the parameter space obtained from Table 1: *ρ* ∈ [0.5 × 10^−3^, 2.5 × 10^−3^] day^-1^,*α*_1_ ∈ [0.01, 1.0] ml/*μ*g day, *α*_2_ ∈ [0.1, 0.75] ml/*μ*g day, *P*(0) ∈ [20, 200], *K* ∈ [300, 550] cm^3^. Virtual trials were run using Matlab 2017b parallel computing toolbox using a parallel algorithm on a 64 GB memory 2.7 GHz 12-core Mac pro workstation under OS X 10.14.

## Results

### The mathematical model describes the general features of LGO response to temozolomide

Typical LGG longitudinal growth and response to therapy consists of four stages (see Fig. 2). First, without treatment tumor grows slowly but steadily [24]. Next, there is an early ‘fast’ tumor volume reduction associated to the start of treatment with TMZ. Finally, after treatment cesation, there is a long-term response. For the patient shown in Fig. 2, the tumor volume reduction lasted for 14 months after the end of the treatment course. Finally, the tumor regrew leading to a clinical relapse. All of those stages were correctly described by the mathematical model. Each stage was associated to one of the biological phenomena reflected as terms in the model equations.

**Fig 2.**
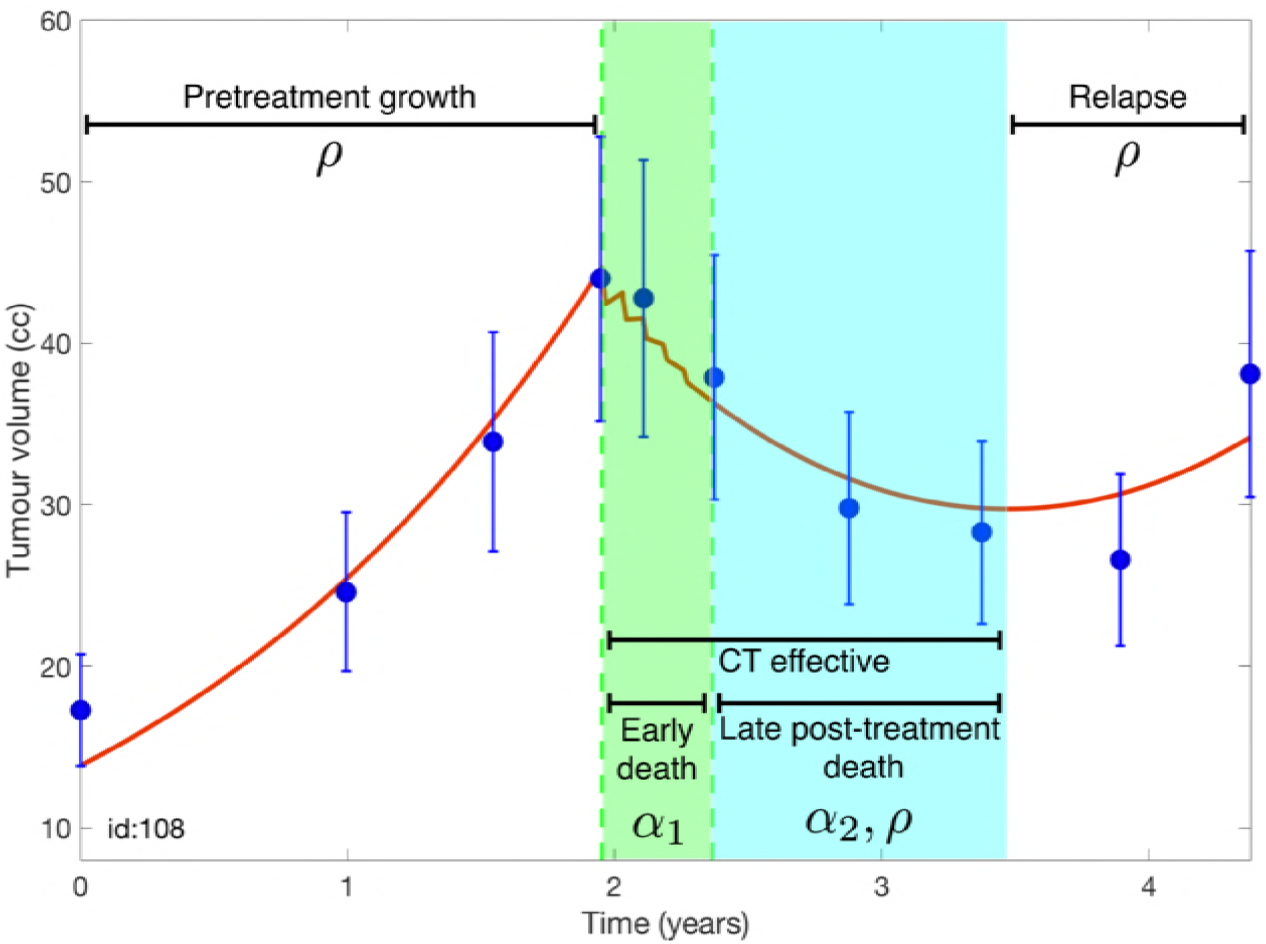
Tumor volumetric longitudinal evolution (solid blue circles) of a patient receiving five chemotherapy cycles together with the best fit found using Eqs. (1) (red lines). Four stages are observed: Pretreatment growth, early response during CT (light green background), post-treatment response (light blue background) and tumor relapse. The model’s parameters modulating the dynamics in each of the stages are also shown.

### The mathematical model describes patient response to temozolomide

We studied the ability of our mathematical model to describe the tumor responses to TMZ. To do so, we fitted the parameters in Eqs. (1) using the longitudinal volumetric data for each patient in our cohort. Figure 3 shows results for selected patients. The model described the longitudinal tumor volumetric data in all cases, what supports the choice of biological mechanisms used to construct it.

**Fig 3.**
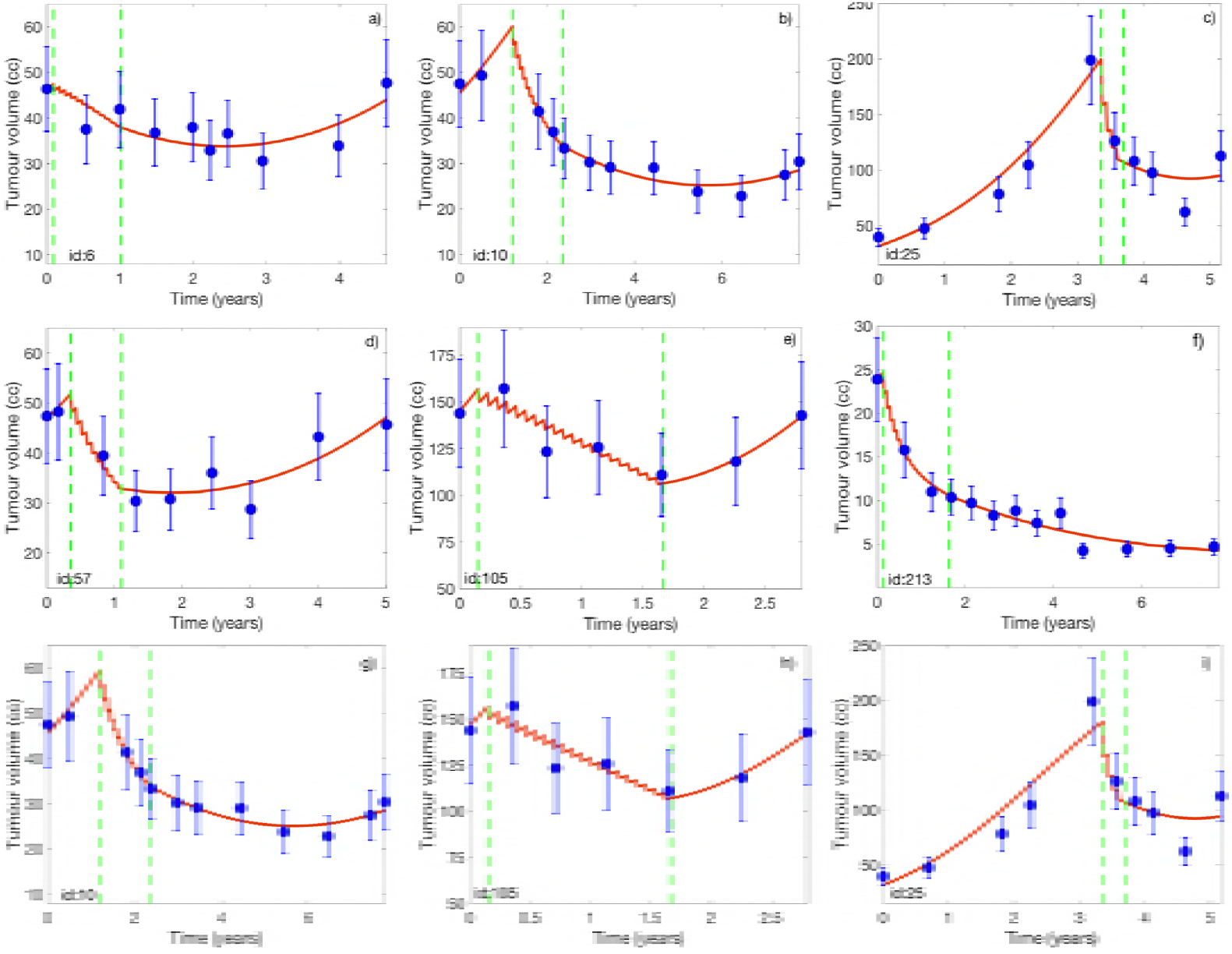
Longitudinal volumetric tumor data (blue circles) and best fits obtained with the model given by Eqs. (1) (red lines). (a-f) Results for six randomly chosen patients from out dataset for a carrying capacity K=523.6 cm^3^. (g-i) Results for three patients for K = 261.8 cm^3^. The vertical dashed lines in each subplots mark the start and end times of treatment with TMZ.

Results shown in Fig. 3 were obtained for a fixed (i.e. non fitted) value of the carrying capacity *K* = 523.6 cm^3^. This parameter provides an estimate of the tumor size for which geometrical and other constraints have a substantial influence on the tumor growth rate. Similar results were obtained for a broad range of values of *K*. As an example, Fig. 3(g-i) shows results for selected patients using a smaller *K* = 261.8 cm^3^. The shapes of the fitting curves and the best root mean square errors (RMSE) were similar for the different *K* values.

### Simulations show potential benefits of alternative treatment schedules

The model was then used as a discovery platform to test alternative treatment regimes in-silico for the patients included in the study. As a first test, we enlarged the time interval between cycles. Five daily doses of the drug were given on days 1-5 of the cycle and then the standard waiting period of 23 days was increased to variable times of up to 6 months. In general, the long-term tumor evolution was similar for all the schedules when the cycle’s length was in the range 1-4 months. Thus, from the volumetric point of view, all schedules led to similar asymptotic dynamics for the in-silico twins of the study patients. Results for selected patients are shown in Figure 4. The only drawback of the long-cycle treatment regimes was the smaller tumor volume reduction observed due to the less intensive nature of the schemes.

**Fig 4.**
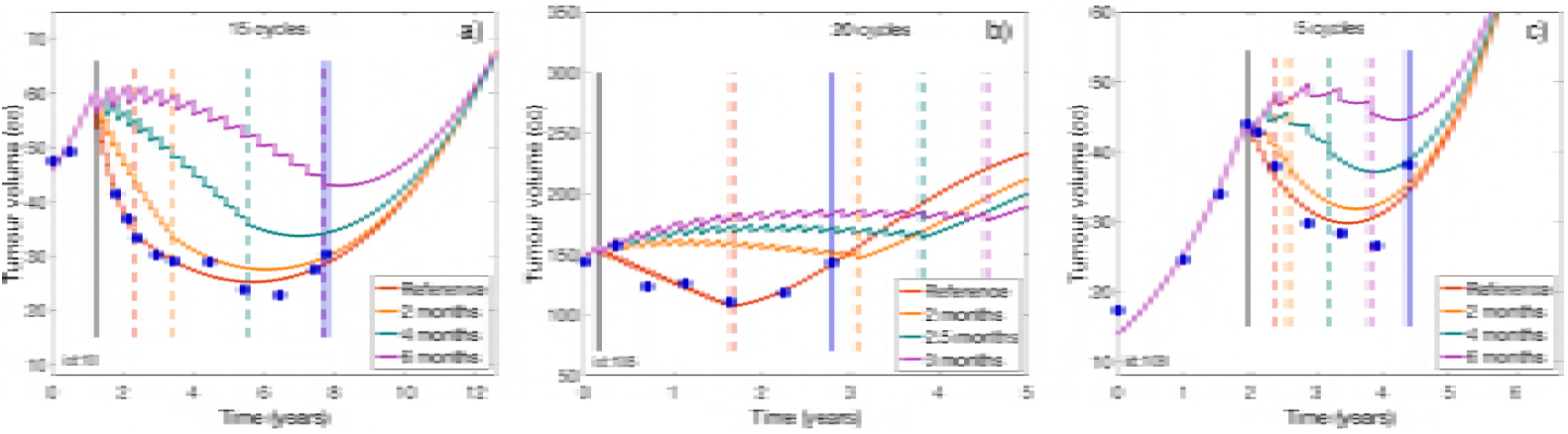
Simulated tumor growth curves under alternative treatment schedules with variable time spacing between consecutive cycles. Esq. (1) were solved for each patient with the best fitted parameters under the different treatment treatment regimens. Each subplot shows the reference (fitted to data) growth curve in red and the simulated growth curves under three alternative schemes with spacing between doses of: (a,c) 2, 4, and 6 months, and (b) 2, 2.5 and 3 months. Vertical dashed lines indicate the end times of the different treatment regimes. The vertical solid blue lines mark the time domain for which imaging follow-up data was available for the patient.

In the case of large tumors, whose size was comparable to *K* and thus the nonlinear term in Eq. (1) played a relevant role, differences between the schemes were observed favoring long-cycle schemes (see e.g. Fig. 4(b)).

As a second series of tests, we explored alternative treatment regimes based on the 28-day cycle. A first regime consisted in 5 doses given following a 1-day on, 1-day off scheme during the first 10 days of the cycle. A second alternative was distributing doses evenly within the cycle duration, i.e. giving a single dose every 4 days. Both treatment regimes led to tumor volumetric evolutions overlapping with the ones of the standard treatment (e.g., those depicted in Figure 3).

Several virtual trials were conducted as described in the ‘Methods’ section. Benefits in median survival were found for the long-cycle strategies that were dependent on the parameter *K*. Long-cycle treatment schemes were never inferior in terms of survival to the standard ones. Indeed, the differences found between survival curves for long-cycle schedules versus the standard ones were never statistically significant (*p* < 0.05) according to the log-rank test.

### A combined treatment regime provided survival advantages in-silico and may provide a standard for LGO patients

Patients in our retrospective dataset were treated with a variable number of TMZ cycles (mean 12, range 4-20, see Table 1). Treatment was effective for all patients included. However, since there is no standardized protocol for chemotherapy in ODG patients, the decision to stop treatment was taken depending on toxicitity, physician and patient preferences, etc.

On the basis of our previous results we explored the potential effectiveness of standardizing treatment for all of the virtual patients consisting of an induction part of five cycles given monthly to reduce substantially the tumor burden followed by a consolidation of 12 cycles given every three months. This treatment scheme was based on the idea that TMZ cycles given every three months should be well tolerated and allow for this long schedule. Moreover, having a first induction part would result in an initial larger tumor volume reduction than for the long-cycle schemes alone. Results are summarized in Figure 5. Survival improvements, many of them substantial, were obtained for the in-silico twins of the patients included in the study (Median 5.69 years, range: 0.67 to 68.45 years, see Fig. 5(a)). Virtual patients for which the number of cycles was larger (patients 3, 6, 7, 8 and 10) than those received by the real one (see Table 1) had larger survival benefits. Also for most patients there was a substantial volumetric reduction in relation to the one achieved for the real patient under the number of cycles given by Table 1 (see Fig. 5(b)).

**Fig 5.**
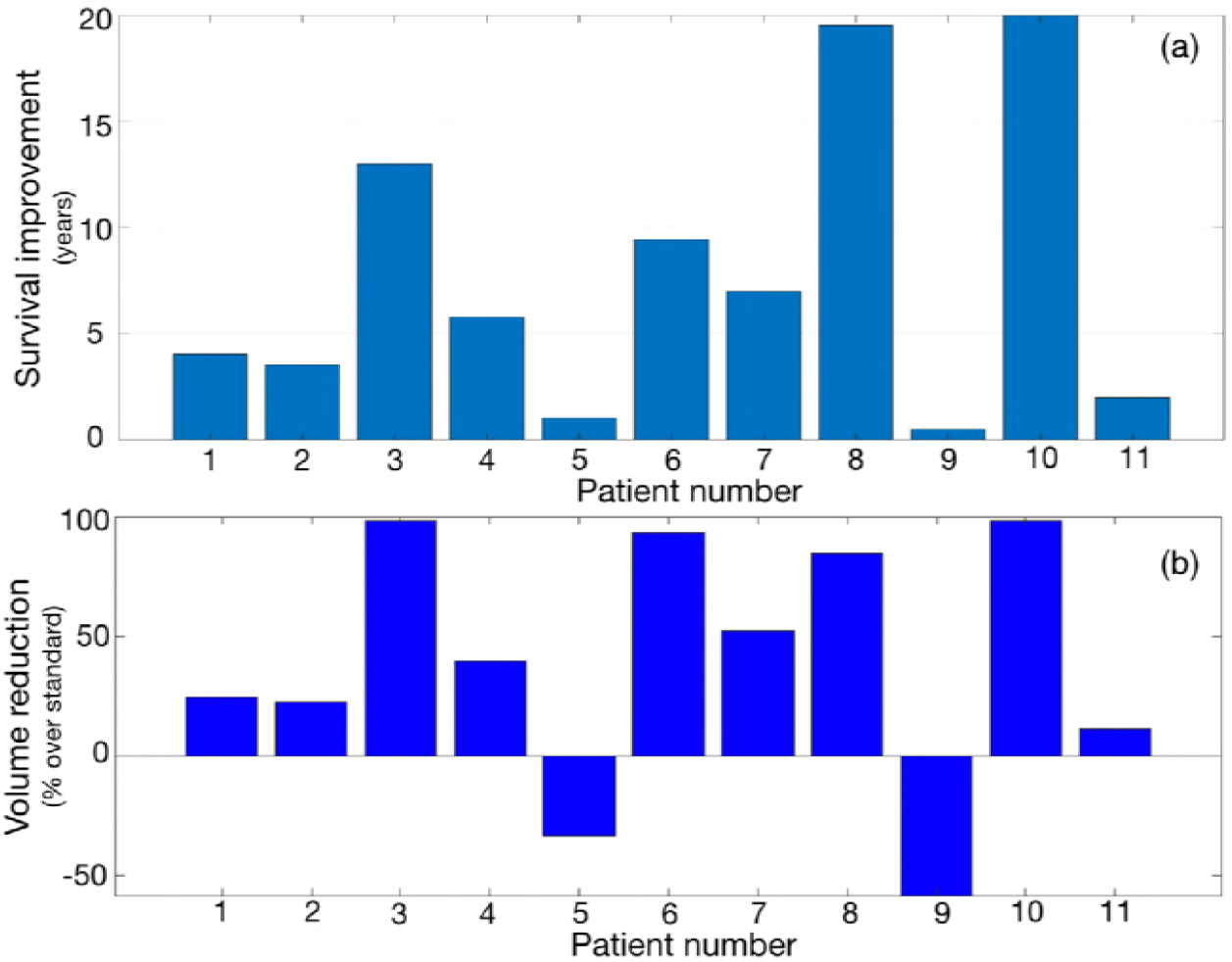
Benefits of the proposed combined treatment with 5 induction cycles given monthly and 12 maintenance cycles given every three months. (a) Predicted survival benefits for the virtual patients subjected to the proposed scheme in comparison with survival of the real patients. (b) Maximum volume reduction obtained by the proposed scheme in comparison with the maximum volume reduction achieved for the real patient.

A virtual trial was run with 2000 virtual patients included in two arms. Results are summarized in Figure 6. Differences between the curves were statistically significant (*p* = 1.65 x 10^−14^, HR = 0.679 (0.614 - 0.75)), with a difference in median survival of 3.8 years between both treatment arms.

**Fig 6.**
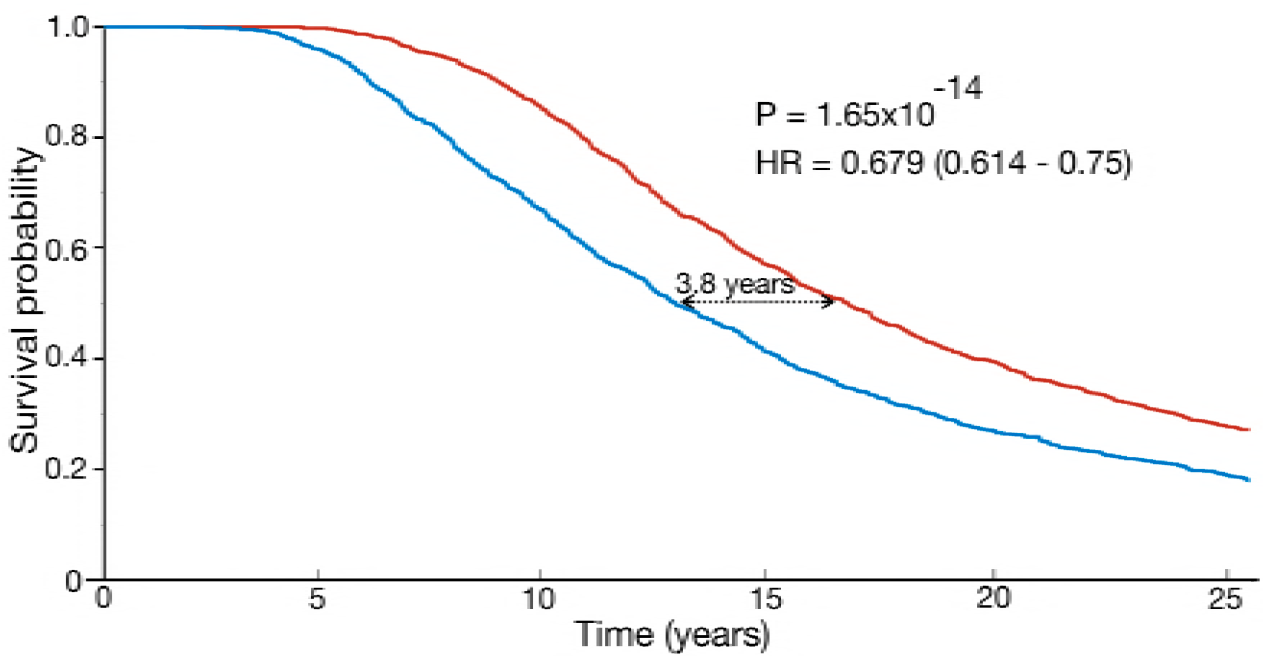
Results of the virtual trial comparing a standard chemotherapeutic approach for LGO versus the proposed scheme. Shownare the Kaplan-Meir plots for both arms. In the first arm (blue), virtual patients received a random number of cycles in the range spanned by the real patients (between 5 and 18 sequential cycles with the standard 1 month spacing). The same virtual patients received the proposed scheme (5 cycles induction given monthly + 12 cycles consolidation given every 3 months). Patients were assumed to die when tumors reached a volume of 280 cm^3^ and those alive after 25 years were considered as censored events.

## Discussion

Our mathematical model successfully reproduced the tumor size dynamics of LGO patients treated with TMZ. That radiological dynamics cannot be described with mathematical models based on instantaneous response to therapy. Thus, a key ingredient in our model was the combination of two different types of death processes, ones leading to ‘early’ cell death (treatment induced apoptosis and necrosis) and others leading to ‘late’ cell death through mitotic catastrophe. This was incorporated through three adjustable parameters. An additional parameter, the carrying capacity, accounts for the limitation of growth due to geometrical constraints. Other parameters were estimated from biological data.

Many authors have built mathematical models to understand and describe different aspects of the natural history and response to treatments of LGGs [25–36], some of them focusing on CT. Ribba et al. [27] developed a six-parameter model based on proliferative, quiescent and damaged quiescent compartments. Some biological assumptions in the model were debatable: there was no connection from the quiescent cell’s compartment to the proliferative cell compartment other than through the damaged quiescent cells, the drug was assumed to affect equally proliferative and quiescent cells, and the drug decay time in brain tissue was fitted to be of the order of several months, a value out of the reasonable range. This is in striking contrast with the value used here inferred from realistic data of a few hours. Bogdanska et al. [34] used a minimal mathematical model incorporating death only through mitotic catastrophe [32], what forced the parameter *κ* to have values beyond the biologically feasible range. Our approach achieved better quantitative fittings than those in Ref. [32] while having all parameter values in meaningful ranges.

For all virtual patients the simulations showed interesting features: (i) Tumor growth was found to be asymptotically similar for different treatment schedules. (ii) There were patients for which a survival increase was observed under the alternative treatment regimes. An obvious implication of (i) and (ii) is that the alternative regimes would have no inferior performance in terms of survival. Our virtual clinical trials also supported those findings.

Those ‘long-cycle’ regimes could have other advantages worth considering. The first one is that they could have substantial benefits in terms of toxicity, pharmacoeconomy and also improving the prognosis. In fact, extending the duration of each cycle is a widely used way to treat toxicities caused by cytotoxics. Another possible benefit of those schemes would be improving drug-exposure in LGOs, by administering a larger number of cycles for longer treatment periods.

The only drawback of long-cycle regimes was a smaller tumor volume reduction due to their less-intensive nature. Although this smaller reduction did not have effects in terms of survival in-silico it would affect symptoms control in real patients. Thus, we designed a mixed treatment scheme consisting of an intensive induction phase of 5 cycles given once per month together with a maintenance stage of 12 cycles given one every three months. This strategy showed an impressive effect on survival. Only for the two patients receiving longer more intensive treatments in real life, the volumetric reductions obtained in-silico were smaller than the ones observed. In spite of that, patients survived longer in the simulations. Indeed, three patients received in real life the same or more CT cycles than in our proposed scheme, but our less toxic scheme resulted in longer survival in the computer simulations. The results were confirmed on a virtual trial including 2000 patients and comparing ‘in-silico’ both treatment arms.

Interestingly, all long-cycle regimes studied were independent on the time point at which the doses were given, i.e. the mathematical model predicts that five doses every 90 days (long-cycle scheduling) would be roughly equivalent to a single dose every 16 days. Thus, choosing one or other regime could be done in terms of toxicity reductions or delaying the appearance of resistant clones. An increasing body of evidence suggests that small subpopulations of cancer cells can evade strong selective drug pressure by entering a ‘persister’ state of negligible growth [37]. This drug-tolerant state has been hypothesized to be part of an initial strategy towards eventual acquisition of bona fide drug-resistance. The induction of persisters in glioma cells has been known to be partially reverted by ‘drug wash-out’ suggesting the contribution of epigenetic mechanisms in drug resistance and supporting the possibility of TMZ rechallenge in glioma patients after prior drug exposure [38], provided there is a sufficiently long waiting time between treatments.

In our work, we assumed a direct proportionality between tumor cell number and the observable tumor size on T2/FLAIR. An interesting extension of this work could be to use partial differential equation-based mathematical models where both quantities are independent. The inclusion of cell-motility processes as in works based on reaction-diffusion models [26,30,31,35] could provide a computational platform to study the delay of the tumor’s malignant transformation through alternative treatment regimes. Further research is required to relate the signal obtained from diffusion MRI sequences and/or ADC maps with local cellularity values.

## Conclusion

We developed a mathematical model of LGOs response to CT describing the longitudinal tumor volumetric dynamics. Once fitted for each patient, the model provided in-silico twins of the real patients. When subjected to long-cycle treatment regimes the ‘virtual twins’ showed similar or better performance in terms of survival. In-silico clinical trials confirmed the results for broader parameter regimes. This long-cycle temozolomide schedules could prove beneficial for LGO patients in terms of toxicity. We studied ‘in-silico’ a treatment combining an induction phase of 5 consecutive cycles plus a maintenance phase (12 cycles given in three-months intervals). The improved drug-exposure of this scheme led to substantial survival improvements and a good tumor control in-silico. We hope this computational study could provide a theoretical ground for the definition of standardized TMZ treatment protocols for ODG patients with improved survival.

## Acknowledgments

We would like to thank Roger Stupp (Northwestern University, USA) for a critical reading of the manuscript and J. Pérez-Beteta (MOLAB, Spain) for his help in the parameter fitting tasks. This research has been supported by the Ministerio de Economía y Competitividad/FEDER, Spain (grant number MTM2015-71200-R), Junta de Comunidades de Castilla-La Mancha (grant number SBPLY/17/180501/000154) and James S. Mc. Donnell Foundation 21st Century Science Initiative in Mathematical and Complex Systems Approaches for Brain Cancer (Collaborative awards 220020560 and 220020450).

